# *Salmonella* infection induces the reorganisation of follicular dendritic cell networks concomitant with the failure to generate germinal centres

**DOI:** 10.1101/2022.12.11.519997

**Authors:** Edith Marcial-Juárez, Marisol Perez-Toledo, Saba Nayar, Elena Pipi, Areej Alshayea, Ruby Persaud, Sian E. Jossi, Rachel Lamerton, Francesca Barone, Ian R. Henderson, Adam F. Cunningham

## Abstract

Germinal centres (GCs) are sites where plasma and memory B cells form to generate high-affinity, Ig class switched antibodies. Specialised stromal cells called follicular dendritic cells (FDCs) are essential for GC formation. During systemic *Salmonella* Typhimurium (STm) infection GCs are absent, whereas extensive extrafollicular switched antibody responses are maintained. The mechanisms that underpin the absence of GC formation are incompletely understood. Here, we show that STm-induces a reversible disruption of niches within the splenic microenvironment, including the T and B cell compartments and the marginal zone. Alongside to these effects post-infection, mature FDC networks are strikingly absent, whereas immature FDC precursors, including marginal sinus pre-FDCs (MadCAM-1+) and perivascular Pre-FDCs (PDGFRβ+) are enriched. As normal FDC networks re-establish, extensive GCs become detectable throughout the spleen. Therefore, the reorganisation of FDC networks and the loss of GC responses are key, parallel features of systemic STm infections.

**Graphical abstract:** 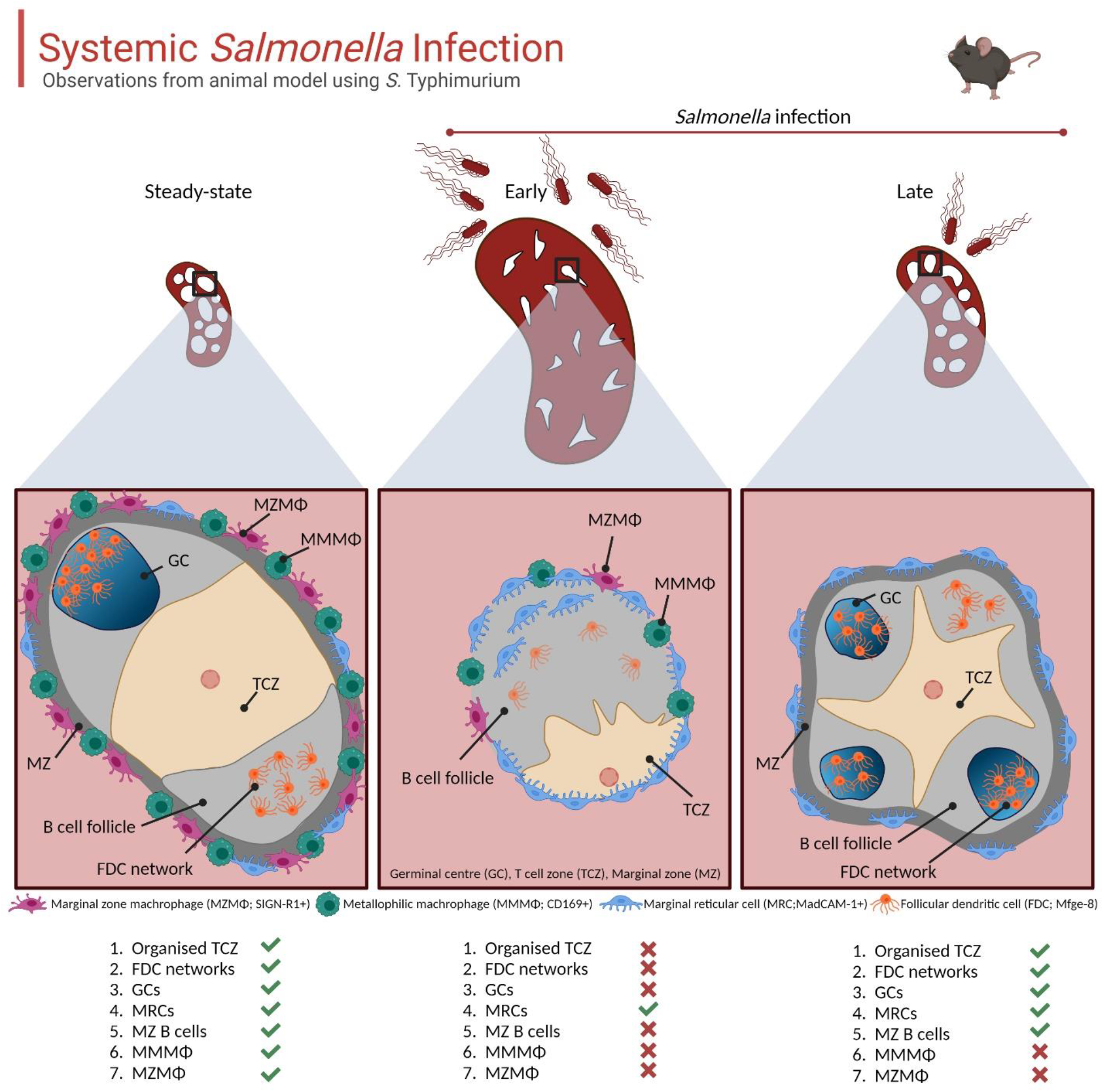

## Introduction

A hallmark of the mammalian immune response is the induction of adaptive immune responses to pathogens. An important contribution to the protection provided by the adaptive immune response is the generation of antibodies, which is a typical consequence of infection and a key aim of vaccination^1^. Antibodies can be generated through two predominant, interconnected pathways. In primary responses, extrafollicular (EF) responses, which develop in the red pulp of the spleen or the medulla in lymph nodes, provide the first wave of IgM and IgG, and these antibodies are typically of modest affinity as there is limited affinity maturation of B cells that enter this pathway^2^. Moreover, antibodies from plasma cells generated through the EF pathway is typically supplanted after a week^3^ or so by antibodies secreted by plasma cells derived from the germinal centre (GC) response^4^. GCs form in the B cell follicles of secondary lymphoid organs (SLO) such as the spleen or lymph nodes^5,6^. A key difference between the EF and GC responses is that antibodies generated from the GC tend to be of higher affinity and most memory B cells and the longest-lived plasma cells derive from this response^7^.

The generation of these productive outputs from the GC requires the interplay of multiple cell types at different sites within SLO, with all processes dependent on the microarchitecture of SLO, including interactions between T and B cells in the T zones and the follicles^8^. Whereas the organisation of cells within T cell zones (TCZ) is dependent upon CCL21/CCL19 secreted by fibroblastic reticular cells (FRC)^9,10^, the generation of B cell follicles relies on CXCL13 secreted by follicular dendritic cells (FDC) and marginal reticular cells (MRCs)^11–13^. Moreover, FDCs play a critical function in the GC by holding antigens in their native conformation on their surface as immune complexes driving the selection of B cell clones with the highest affinity^14,15^. Therefore, the organisation of lymphoid tissues is essential for the efficient generation of productive immune responses and FDCs are crucial for normal follicle architecture and the GC reaction^16,17^.

Antibody responses induced during natural infection can help moderate the spread of the pathogen and secondary superinfections. Nevertheless, many pathogens can modulate the capacity to mount antibody responses during infection, potentially affecting the capacity of the host to deal with current or later infectious threats. Bacterial and parasitic infections such as those caused by *Ehrlichia muris, Salmonella* Typhimurium (STm) and *Plasmodium* sp. induce atypical GC responses^18–21^. Indeed, in models of STm infection, GCs are not detectable until a month after infection, whereafter there is a significant increase in serum antibody titres, and these are of a higher affinity ^20,22,23^. The altered kinetics described for these pathogens are markedly different to the kinetics and processes described for GC responses that develop to non-viable, proteinaceous T-dependent antigens such as alum-precipitated ovalbumin or chicken gamma globulin. In these models, the GC response is established by a week after immunization, and the process is complete by around five weeks after immunization^24^. Moreover, pathogens such as STm can impair the induction of GC responses to co-immunized antigens, demonstrating that this effect is not restricted to bacteria-associated antigens ^25–27^. The reasons for this delay in GC induction are incompletely understood but is not due to a failure to induce STm-specific T and B cell responses. For instance, within 24 hours of STm infection, T cell priming and Th1 cell differentiation is already established, and extensive EF antibody responses are detectable soon after this^20,26^. Despite these gaps in our understanding, some insights into why STm has these effects have been reported. Factors that influence the level of the GC response include the number of viable bacteria present^20,23^, reduced T follicular helper (Tfh) cell differentiation by IL-12 mediated induction of T-bet^28^, the recruitment of Sca-1+ monocytes to lymphoid organs and ineffective cellular respiration dependent on TNF and IFNγ^27^. Despite these insights, it is unclear whether STm infection influences stromal cell populations and the compartments within SLO after infection. Here, we assessed the topological changes in the spleen and investigated the role of FDCs in the lack of GCs during acute STm infection and found that STm infection can modulate mature FDC networks for weeks and only when FDC networks reform do GC develop.

## Methods

### Mouse and bacteria strains and infection of mice

All the experiments were performed following the animal research regulations established by the Animals (Scientific Procedures) Act 1986 (ASPA) and under the licenses, P06779746 and I01581970 issued by the Home Office. Wild-type adult C57BL/6 mice (6-8 weeks) were purchased from Charles River Laboratory and housed in the Biomedical Services Unit at the University of Birmingham during the infection period. Mice were systemically infected (intraperitoneally) with 5 × 10^4^ colony-forming units (CFU) of aroA-deficient *S*. Typhimurium SL3261 attenuated strain as previously described^20,42^. Control mice were injected with sterile-filtered Dulbecco’s-phosphate-buffered saline (DPBS). Spleens were collected at the indicated time points after the infection. Bacteria load per organ was quantified by plating 10-fold diluted tissue homogenates on Luria-Bertani agar plates.

### Flow cytometry

Single-cell suspensions were obtained by mechanical disaggregation of the spleen across 50 μm-cell strainers and washed with cold RPMI-1640 medium containing 5% fetal bovine serum (FBS) and 5 mM ethylenediaminetetraacetic acid (EDTA). Red blood cells were lysed using ammonium-chloride-potassium (ACK) lysing buffer (Life Technologies). Cells suspensions were incubated with a CD16/32 antibody (eBioscience) in FACS buffer (2% FBS, 5mM EDTA and 0.01% of sodium azide in PBS) to reduce FcγRIII/FcγRII-mediated non-specific binding. Fixable viability dye eFluor 450 (eBioscience) was used to discriminate between live and dead cells. Cell suspensions were incubated with a mix of the following antibodies in FACS buffer: anti-B220 (RA3-6B2; BD Bioscience), anti-CD19 (1D3; BD Bioscience), anti-IgD (11-26c.2a; BioLegend), anti-IgM (II/41; eBioscience), anti-CD21/35 (7E9; BioLegend) and anti-CD23 (B3B4; BD Bioscience). To quantify stromal populations, a portion of spleens was digested on a solution containing 0.2 mg ml^−1^ collagenase P (Roche), 0.1 mg ml^−1^ DNAase I in RPMI-1640 medium and enriched by depleting CD45+ cell and TER119+ cells using MACS microbeads (Miltenyi Biotec). Cells suspensions were incubated with anti-CD45 (30-F11; eBioscience), anti-erythroid cells (TER-119; BD Bioscience), anti-Podoplanin (8.1.1; eBioscience), anti-CD31 (390; BD Bioscience), anti-MadCAM-1 (MECA-367; BD Bioscience) and anti-CD21/35 (7E9; BioLegend). Samples were acquired using a BD LSR II Fortessa flow cytometer and data was analysed using Flowjo Software v10.8.1 for Windows (Ashland, OR, USA).

### Immunohistochemistry

Spleens were obtained at different time points after infection as indicated in each figure, and frozen in liquid nitrogen. Five-μm slices were obtained using a Bright cryostat and fixed with acetone at 4°C for 20 min. Slides were rehydrated in Tris Buffer (TBS; pH 7.6) and labelled with the following antibodies for immunohistochemistry analysis: anti-CD3 (145-2C11; eBioscience), anti-IgD (11-26C.1; BD Bioscience), anti-follicular dendritic cell (FDC-M1; BD Bioscience) and MadCAM-1 (MECA-367; eBioscience). The following secondary antibodies were used and incubated at room temperature for an hour: Peroxidase-conjugated goat anti-Armenian Hamster IgG (H + L) from Fitzgerald, Biotin-SP-AffiniPure Donkey Anti-Rat IgG (H+L) and Peroxidase-AffiniPure Donkey Anti-Rat IgG (H+L) from Jackson ImmunoResearch (Cambridge, UK). Peroxidase activity was detected using SIGMAFAST3,3’-Diaminobenzidine (DAB) tablets and hydrogen peroxide dissolved in TBS buffer pH 7.6. Alkaline phosphatase enzyme activity was detected using VECTASTAIN[R] ABC-AP Kit (Vector Laboratory) and developed with a mix of Naphthol AS-MX Phosphate-free acid, fast blue B salt and Levamisole [(-)-tetramisole hydrochloride] in TBS buffer pH 9.2. All chemical reagents for immunohistochemistry were purchased from Sigma-Aldrich. Slides were mounted in VectaMount Permanent Mounting Medium (Vector Laboratories), and the total spleen area was imaged using a microscope slide scanner (Zeiss Axio Scan.Z1).

### Immunofluorescence microscopy

Cryosections were rehydrated in PBS-0.01% Tween 20 and incubated at room temperature with the primary antibodies in a PBS solution containing 1% of bovine serum albumin, 1% of normal human serum and 0.01% of sodium azide. The following antibodies were used: anti-IgD (11-26C.1; BD Bioscience), anti-B220 (RA3-6B2), anti-CD3 (145-2C11; eBioscience), anti-laminin (Polyclonal; Invitrogen), anti-CD209b (SIGN-R1; 22D1; eBioscience), anti-CD169 (Siglec-1; 3D6.112; BioLegend), anti-CD205 (205yekta; eBioscience), anti-CD11c (N418; BioLegend), anti-IgM (Polyclonal; Southern Biotech), anti-milk fat globule epidermal growth factor 8 protein (Mfge-8; 18A2-G10; MBL), anti-fibroblasts monoclonal antibody (ER-TR-7; Invitrogen), anti-follicular dendritic cell (FDC-M1; BD Bioscience), anti-CD21/35 (REA800; Miltenyi Biotec), anti-CD31 (Polyclonal; R&D System), anti-PDGFRβ (APB5; eBioscience), anti-CCL21 (Polyclonal; R&D System) and anti-CXCL13 (Polyclonal; R&D System), anti-MadCAM-1 (MECA-367; eBioscience), anti-Ki-67 (Polyclonal; Abcam), peanut agglutinin (PNA; Vector Laboratories), and *anti-Salmonella* (Polyclonal; Abcam). The following fluorescent-labelled streptavidin and secondary antibodies were used to detect unconjugated antibodies and incubated for 45 min in the dark: AlexaFluor 488-conjugated anti-rat IgG, Cy3-conjugated anti-rat IgG, AlexaFluor 647-conjugated anti-hamster IgG, Cy3-conjugated anti-rabbit IgG, AlexaFluor 555-conjugated streptavidin, AlexaFluor 488-conjugated streptavidin, Brilliant Violet 421-conjugated streptavidin. Four-step indirect IF staining was performed for the detection of chemokines using AlexaFluor 488-conjugated anti-sheep IgG, rabbit anti-AlexaFluor 488, and AlexaFluor 488-conjugated anti-rabbit IgG. Slides were mounted with ProLong Glass Antifade Mountant (Thermofisher Scientific), and images were acquired with Zeiss Axio Scan.Z1 or Zeiss LSM780 Confocal Microscope. Pixel intensity quantification was performed with Fiji (ImageJ) software (National Institutes of Health) and ZEN 3.0 Software.

### Laser Capture Microdissection, RNA isolation and quantitative PCR

Eight-micron sections for microdissection were cut onto membrane slides (Carl Zeiss™ 1.0 PEN NF 41590-9081-000), stained with 1% (v/v) cresyl violet acetate and stored at −80°C. Slides were bought to room temperature and microdissection was performed using the PALM Robo Software V.4.6 software using the brightfield “AxioCam CC1” setting on a Zeiss PALM MicroBeam Laser Capture Microdissection microscope. Microdissected tissue was collected into tubes containing 20 μl of RLT buffer and β-Mercaptoethanol. Afterwards, collection tubes were stored at −80°C prior to RNA extraction. RNA extraction from the microdissected samples was completed as per the protocols provided using the RNeasy FFPE kit (Qiagen). The RNA was then reverse transcribed using the high-capacity reverse transcription cDNA synthesis kit (Applied Biosystems) according to the manufacturer’s specifications. Quantitative RT-PCR (Applied Biosystems) was performed on cDNA samples for *Ccl19, Cxcl13, Lta, Ltb, Ltbr, Tnfr1* and *Tnfa* mRNA expression. β-actin was used as an endogenous control. The primers and probes used were from Applied Biosystems, and samples were run in duplicates on a 384-well PCR plate (Applied Biosystems) and detected using an ABI PRISM 7900HT instrument. Results were analysed with the Applied Biosystems SDS software (SDS 2.3). The mean of two technical replicates (C_t_ values) was used to calculate the ΔC_t_ value for which the C_t_ of the β-actin was subtracted from the C_t_ of the target gene C_t_ value, and the relative amount was calculated as 2^−ΔC^t. C_t_ values above 34 were not accepted, and neither were technical replicates with more than two-cycle differences between them.

### Statistical analysis

Statistical analysis was performed using GraphPad Prism 9 for Windows (Version 9.4.1 (681). The bar height in all graphs represents the median, and the error bars display the 95% confidence interval (CI). Two-tailed, unpaired, *t*-test was applied when comparing two groups, and one-way ANOVA and Tukey’s or Dunnett’s multiple comparison test was performed when comparing between groups or against the non-infected (NI) control, respectively. A significant difference between groups was considered when the *p*-value was < 0.05.

## Results

### *Salmonella* Typhimurium infection perturbs the organisation of the white pulp microarchitecture

Systemic infection of susceptible C57BL/6 mice with attenuated STm SL3261 results in a self-resolving infection characterised by rapid colonisation of the spleen and bacterial numbers peaking from the first week before gradually decreasing after the third week (Fig. 1a). In parallel, STm infection induces a marked splenomegaly which peaked at 21 days, when spleens were around 10 times the mass of non-infected control mice (Fig. 1b). As reported previously^20^, GCs are not a feature of early STm infections and are only consistently detected at day 42 after infection when the infection and associated splenomegaly has largely resolved (Fig. 1c-d). We hypothesised that GC responses are delayed during STm infection because infection induces a perturbed white pulp (WP) topography thus inhibiting the generation of productive responses. Assessment of the WP containing the combined T and B cell compartments in the spleen showed that the proportion of the spleen that is WP decreased between 7 and 30 days after infection before recovering to similar proportions as non-infected mice by day 42 post-infection (Fig. 2a, b). Moreover, at day 21, when the effects of infection on splenic architecture are most pronounced, the absolute area of individual white pulps was significantly smaller than in control mice and contained poorly defined T and B cell areas, with a relative paucity of T and B cells within their respective compartments (Fig. 2a-e). Features that characterised the WP post-infection included the altered distribution of dendritic cells (DCs; DEC205+ CD11c+ cells). These cells are mostly restricted to the TCZ in non-infected mice, but were found throughout the WP area after infection, including in and around B cell follicles (Fig. 2f; Supplementary Fig. 1a).

**Figure 1.**
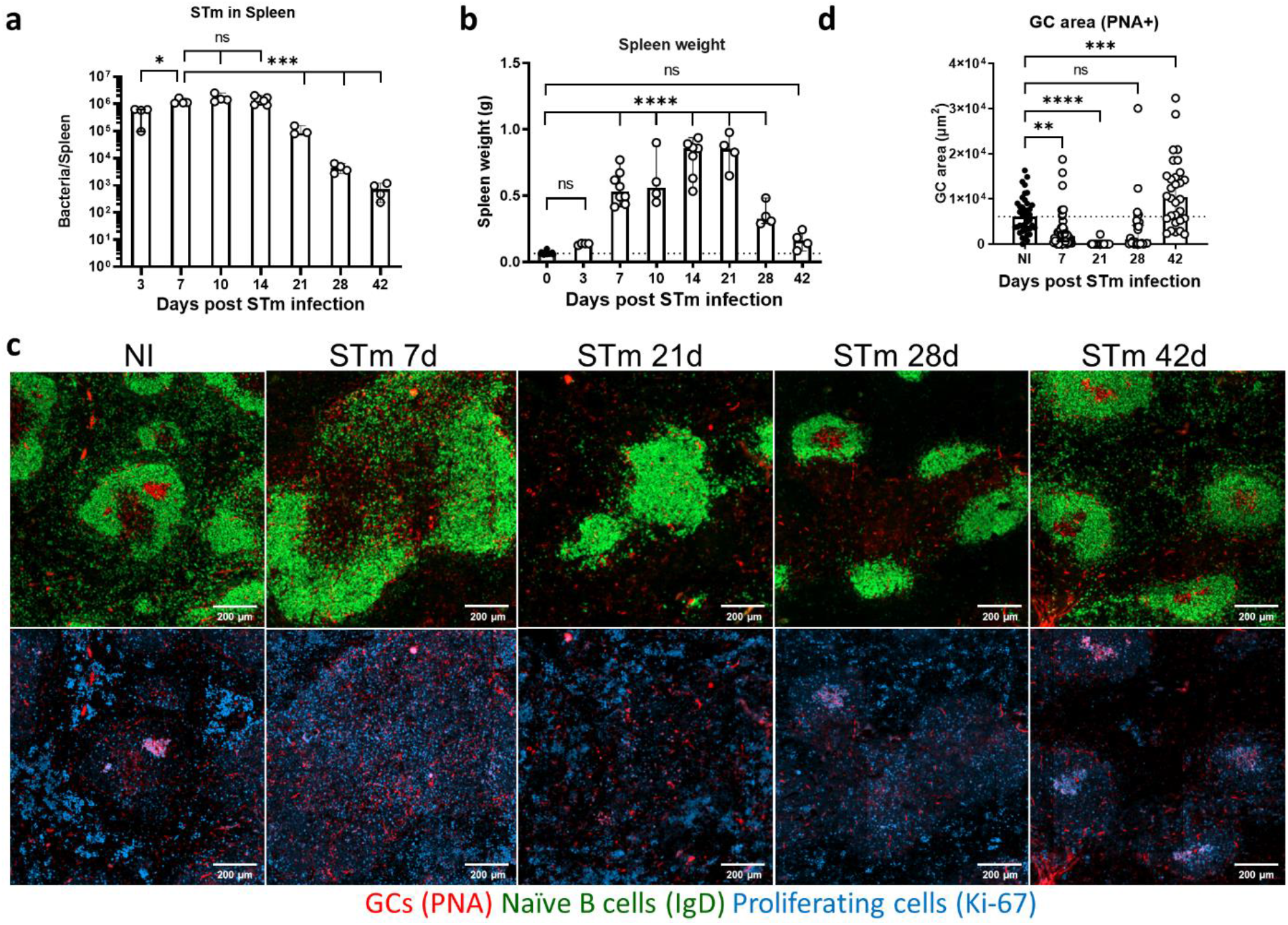
Kinetics of GC development during STm infection. Mice were infected i.p with 5×10^5^ STm SL3261 and the spleens recovered at the indicated times. **a** Bacteria loads (CFU) per spleen. Each symbol represents one mouse. **b** Spleen weights from individual infected and NI mice. **c** Top row: representative images of spleen sections from infected and NI mice stained to detect IgD expressing cells (green) and PNA binding cells (red); bottom row: the same area as per top row showing PNA-binding and Ki-67 expressing cells. Scale bars 200 μm. **d** Quantification of GC area determined by measuring 45 follicles per section. Each point represents the area measured for each GC. Data is representative of 2-3 experiments with 4 mice each, the bar height represents the median, and the error bars display the 95% CI. One-way ANOVA and Tukey’s multiple comparison test for **a** or Dunnett’s multiple comparison test for **b** and **d** were performed. \**p* < 0.05, \*\**p* < 0.01, \*\*\**p* < 0.001, \*\*\*\**p* < 0.0001. ns, nonsignificant.

**Figure 2.**
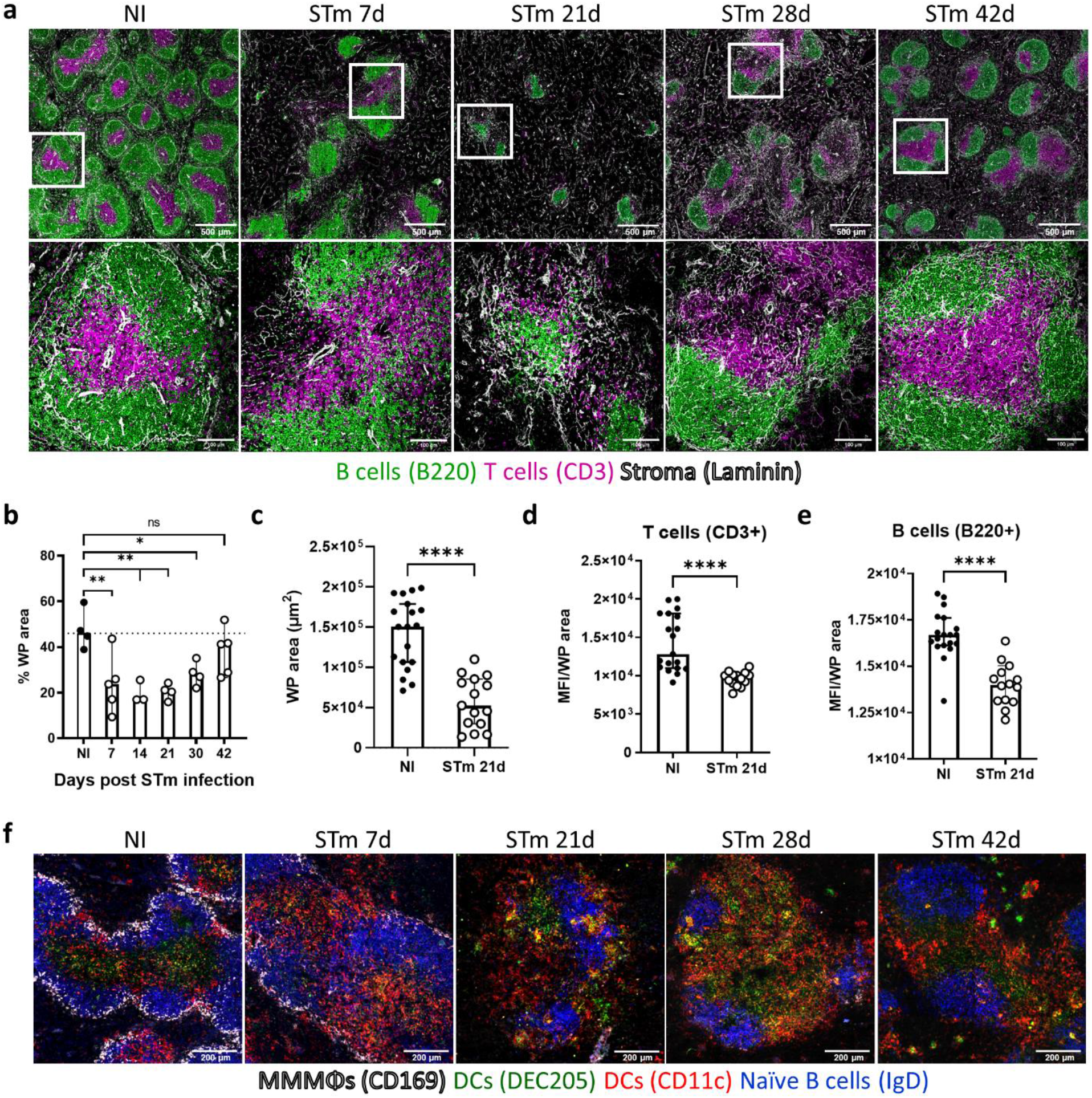
STm infection-induced alteration of the splenic microarchitecture. Mice were infected as per Fig. 1. **a** Cryosections from spleens were stained to detect T cells (CD3+; magenta), B cells (B220+; green), and the reticular fibre network (basement membrane; laminin+; white) to define the compartments in the WP. Representative low-magnification images are shown in the top row (scale bar 500 μm), and confocal images in the bottom row represent higher magnifications of the selected areas. Scale bar 100 μm. **b** Graph showing the proportion of WP per total area of spleen section. Each symbol represents a single mouse. **c** Graph showing the area (in μm^2^) of 15 randomly selected individual WP. **d**, **e** The density of B and T cell staining respectively per WP (determined as the median fluorescence intensity (MFI) of the signal for B220 and CD3). **f** Cryosections from spleens were stained to detect MMMΦs (CD169+; white), DCs (DEC205+ in green and CD11c+ in red), and naïve B cells (IgD+; blue). Scale bar 200 μm. Images and *in situ* quantification representative of 2 experiments with at least 4 mice in each group, the bar height represents the median, and the error bars display the 95% CI. One-way ANOVA and Dunnett’s multiple comparison test for **b**, and two-tailed, unpaired, *t*-test for **c-e** was performed. \**p* < 0.05, \*\**p* < 0.01, \*\*\*\**p* < 0.0001. ns, nonsignificant.

The marginal zone (MZ) borders the WP and discrete populations of macrophages and B cells reside in this site, including CD169+ metallophilic macrophages (MMMΦs) and SIGN-R1+ MZ macrophages (MZMΦs) as well as MZ B cells. Moreover, the cell populations in the MZ can regulate GC B cell responses^29,30^. By day 7 after infection, immunofluorescence (IF) microscopy showed a reduced detection of CD169+ MMMΦs, which was more apparent from day 21 and afterwards, and a near absence of signal for SIGN-R1+ MZMΦs (Fig. 3a-c; Supplementary Fig. 1b). Additionally, STm infection induced a reduction in B cells in the MZ as assessed by both imaging and flow cytometry (Fig. 3a, d-f). Whilst MZ B cells recovered by day 42, this was not the case for CD169+ MMMΦs and SIGN-R1+ MZMΦs. Therefore, STm infection results in a significant remodelling of the WP and MZ splenic microarchitecture.

**Figure 3.**
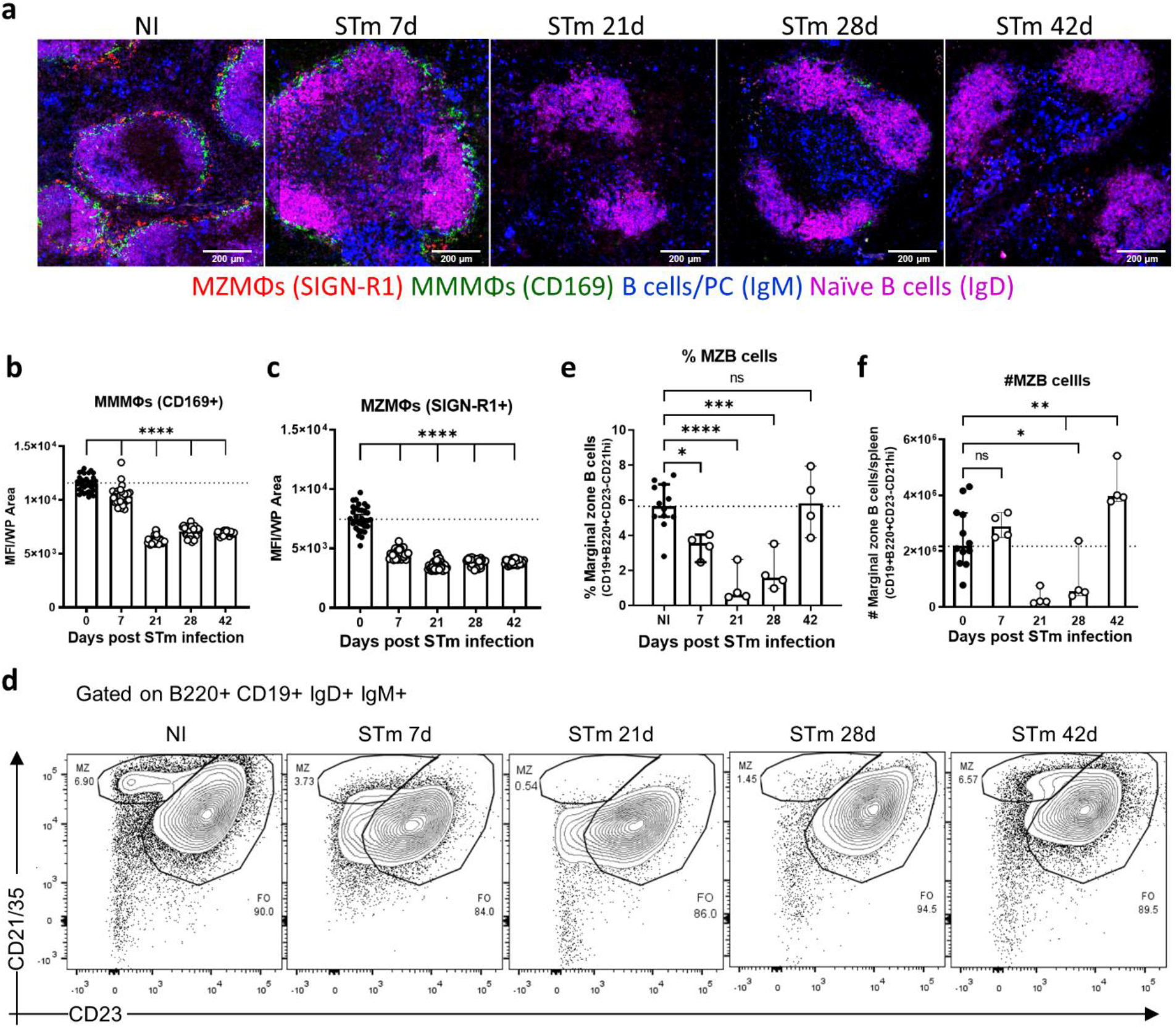
STm infection-induces changes in macrophages and B cells in the MZ. Mice were infected as per Fig. 1. **a** Cryosections from spleens were stained to detect MMMΦs (CD169+; green), MZMΦs (SIGN-R1+; red) and MZ B cells (IgM+; blue). Scale bar 200 μm. **b, c** *In situ* quantification of the signal for CD169 and SIGN-R1, respectively, in individual WP was measured and expressed as MFI. Each symbol represents the MFI per WP. **d** Representative flow cytometry contour plots for the identification of MZ B cells. MZ B cells were gated on B220^+^CD19^+^IgM^+^IgD^+^CD21^hi^CD23^int^ expression. **e, f** Graphs displaying the percentage and absolute number, respectively, of MZ B cells at the indicated times. Data is representative of 2 experiments with at least 4 mice each, the bar height represents the median, and the error bars display the 95% CI. One-way ANOVA and Dunnett’s multiple comparison test was performed for **b, c, e** and **f**. \**p* < 0.05, \*\**p* < 0.01, \*\*\**p* < 0.001, \*\*\*\**p* < 0.0001. ns, nonsignificant.

### STm-induced loss of organised FDC networks correlates with the lack of GCs

The perturbed microarchitecture observed after infection suggested that stromal cells which orchestrate the migration of cells in SLO might be affected by STm. FDCs play a critical role in the antigen-mediated selection of B cell clones in the GC, and since organised GCs are not detected in the first weeks after infection, we examined the localisation of FDCs networks after infection. FDCs can be identified by the expression of milk fat globule epidermal growth factor 8 protein (Mfge-8) in conjunction with the expression of CD21/35. The classic anti-FDC antibody FDC-M1 recognises Mfge-8^31^. Spleens from non-infected and infected mice were stained with the FDC-M1 antibody to detect FDCs (Fig. 4a). This identified a classical distribution of FDC networks in follicles and immature FDCs in the MZ in non-infected mice. Seven days after STm infection, there was a noticeable decrease in the number of compact FDC networks, instead, a filiform reticular pattern staining in the MZ predominated. By day 14 some FDC-M1+ cells were also found as individual cells distributed throughout the WP. At 21 days, few FDC-M1+ cells were detected within the follicles. Flow cytometry showed that there was a reduced density, but not absolute number of FDCs in spleens at day 21 post-infection (Fig. 4b-d), reflecting the significant increase in spleen size (Fig. 1b) and cellularity at this time^32,33^.From day 28 post-infection classical FDC networks started to organise again, and these appeared normal by day 42 (Fig. 4a-d; Supplementary Fig. 2a). These findings were confirmed using a different IF staining panel with spleen sections co-stained for complement receptors (CR1/2; CD21/35) which are highly expressed in FDCs and with anti-Mfge-8 (18A2-G10), which target different epitopes than FDC-M1 (4C11)^31^ (Fig. 4e; Supplementary Fig. 2b). This confirmed the reorganisation of the FDC networks and the *in situ* quantification showed a reduced MFI of FDC markers in WP (Supplementary Fig. 3a, b). This was not related with direct infection of FDC or follicles by STm as few bacteria were detected in follicles or associated with FDC (Supplementary Fig. 3c). Next, the lack of organised FDC networks and the development of GCs were evaluated in parallel. Spleen sections from non-infected mice and STm-infected mice stained with peanut agglutinin (PNA), anti-IgD, and anti-Mfge-8. GCs (PNA+ IgD-) were only detected when classical FDC networks were recovered, for instance at 28 days few GCs were detected only in follicles displaying a more compact FDC network and by 42 days, nearly all follicles contain a GC and an organised FDC network (Fig. 5a). The maintenance of the FDC network in the adult spleen is dependent on LT/LTβR and TNF/TNFR signalling^34–36^. Gene expression of *Ltb, Lta, Ltbr, Tnf*, and *Tnfr1* was investigated by RT-PCR in microdissected WP isolated from the spleens of non-infected mice and mice infected for 21 days. *Ltb* expression was significantly downregulated (Fig. 5a), whereas *Ltbr and Tnfr* gene expression was significantly upregulated in WP of infected mice compared to non-infected mice (Fig. 5c, d). *Tnf* and *Lta* expression was not different between the groups (Fig. 5e, f). Overall, the transient absence of GC is associated with the lack of FDC networks and perturbations in gene expression in the lymphotoxin and TNF pathways.

**Figure 4.**
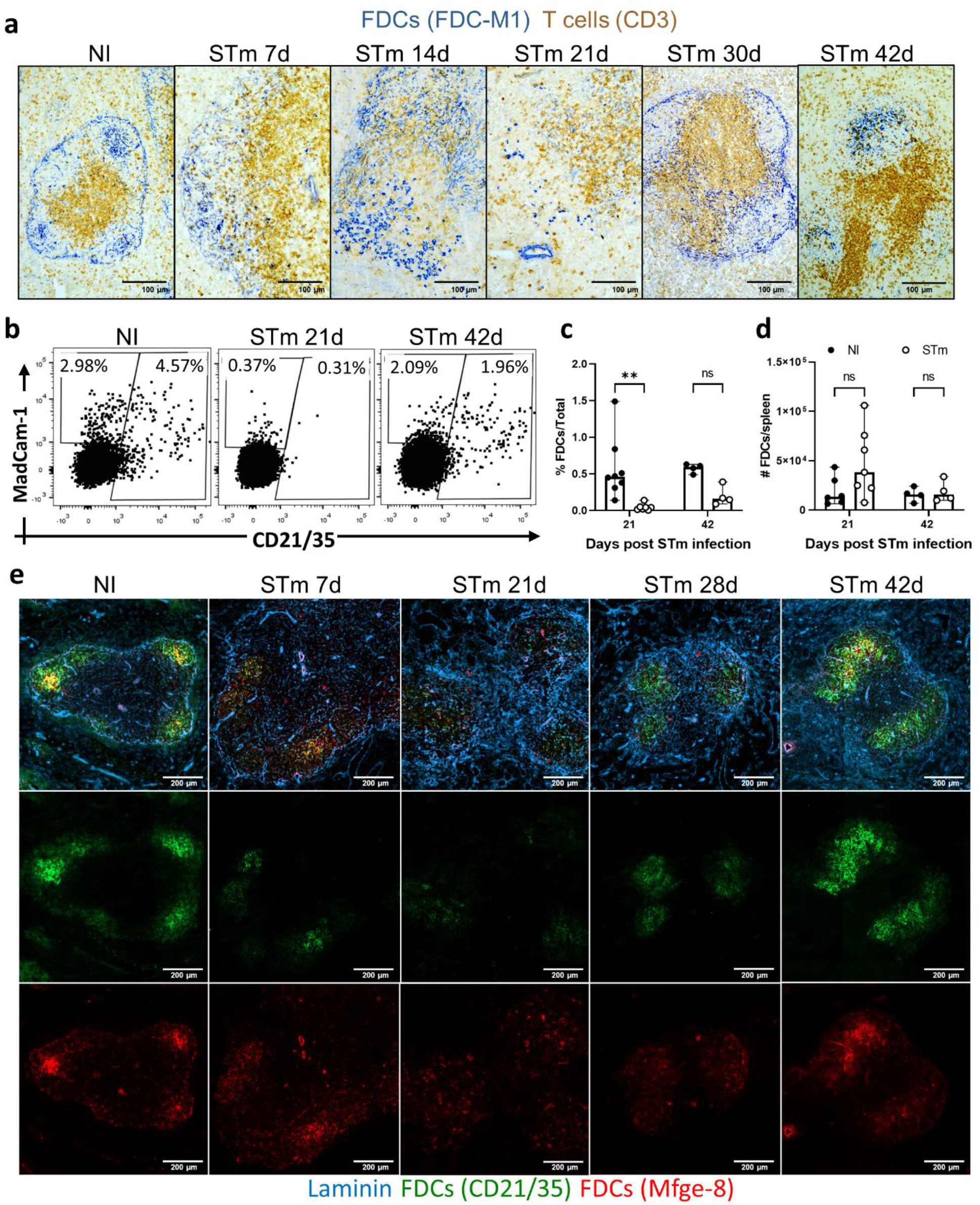
FDC network remodelling during STm infection. Mice were infected as per Fig. 1. **a** Representative images of spleen cryosections stained by immunohistochemistry to detect FDC (FDC-M1+; blue) and T cells (CD3+; brown), scale bar 100 μm. **b** Representative dot plots of FDCs and MRCs detected in spleens of NI mice and mice infected for 21 and 42 days. Cells were gated on non-haematopoietic cells (CD45), non-endothelial (CD31), non-erythroid (TER119) cells and based on the expression of podoplanin, MadCAM-1 and CD21/35. **c, d** Graphs showing the proportion of total cells and absolute number of FDCs at 21 days and 42 days after STm infection. Each point represents results from one spleen, the bar height represents the median, and the error bars display the 95% CI. Two-way ANOVA and Šídák’s multiple comparison test was performed for **c-d**. \*\**p* < 0.01; ns, nonsignificant. **e** IF images stained to detect laminin (blue), FDCs (Mfge-8+; red), and CR1/CR2 (CD21/35+; green). Top row shows a merged image of the markers, and the middle and bottom rows show single-colour images of the same area. Scale bar 200 μm.

**Figure 5.**
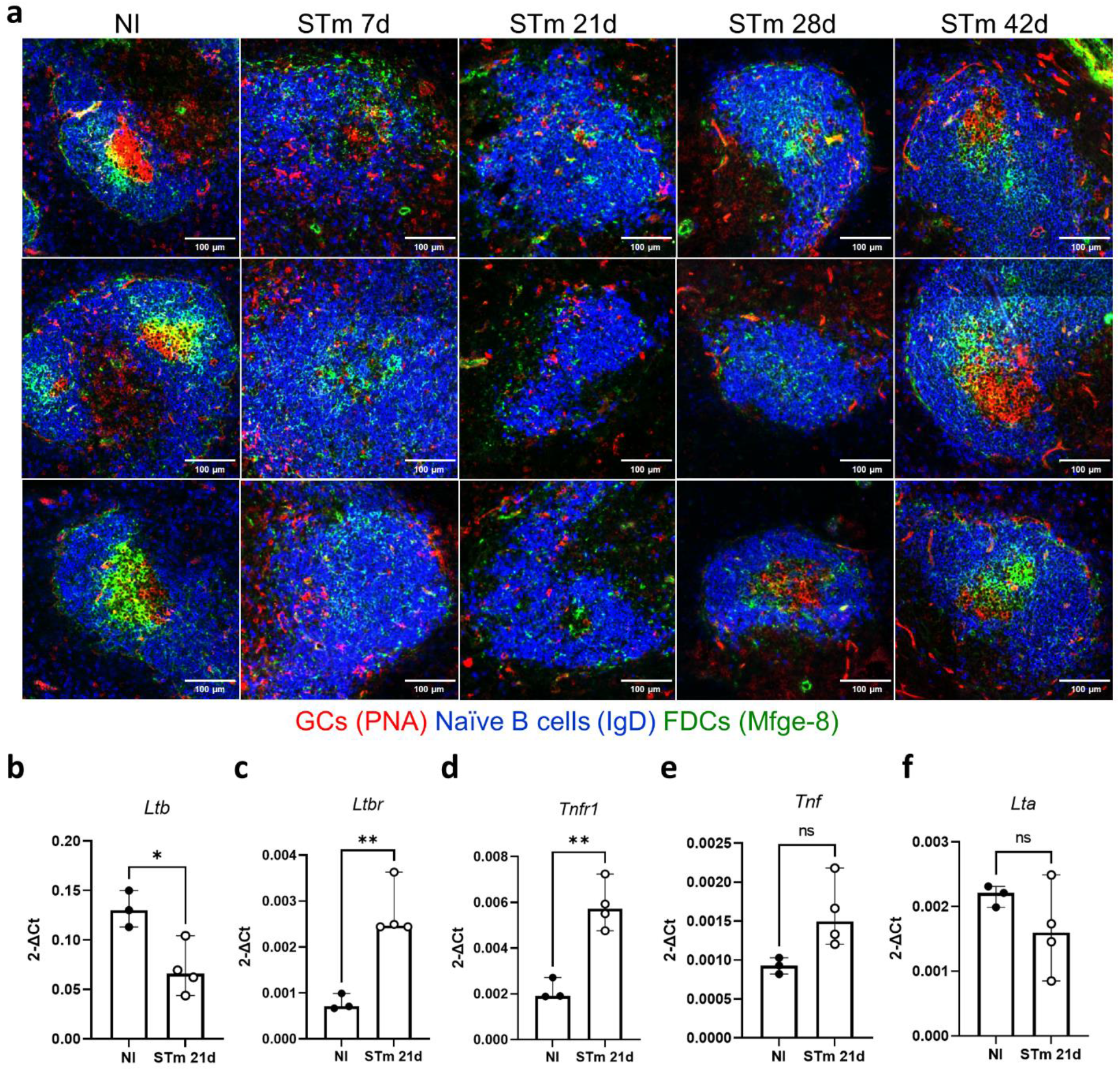
GCs are detected in parallel with the normalisation of FDC networks. Mice were infected as per Fig. 1. **a** Representative IF images of B cell follicles (IgD+; blue), FDCs (Mfge-8+; green), and GCs (PNA+; red) detected in spleens of infected and NI mice. Scale bar 100 μm. **b-f** Graphs showing the expression of individual genes in WP isolated by microdissection from NI mice or mice infected for 21 days. Each point represents the gene expression detected in WP from one individual mouse, the bar height represents the median, and the error bars display the 95% CI. Two-tailed unpaired, *t*-test was used to compare groups. \**p* < 0.05, \*\**p* < 0.01, ns, nonsignificant.

### Perivascular and marginal sinus FDC precursors expand during infection

We assessed the effects of infection on other stromal cells that may contribute to follicle organisation. One such cell-type are MRCs, which express the adhesion molecule MadCAM-1, secrete CXCL13 and have been described as marginal sinus pre-FDCs as they also share the expression of Mfge-8 in the MZ in steady-state^13,37^. Spleen sections from non-infected mice or mice infected with STm were stained for MadCAM-1, anti-Mfge-8 and anti-CD21/35 and assessed by IF (Fig. 6a; presented as a merge of all staining on the top row or just Mfge-8 and MadCAM-1 on the bottom row). STm infection induced a noticeable expansion of MadCAM-1+ cells from 7 days up to 28 days after the infection, and these cells were detected not only in the MZ but also in the follicles, and some were positive for Mfge-8 (Fig. 6a; Supplementary Fig. 3d). Analysis by flow cytometry confirmed that there was an expansion in the frequency and number of MRCs at day 21 post-infection, which was not observed at day 42 (Fig. 6b). Additionally, FDC-M1+/Mfge-8+ cells were observed around CD31+ blood vessels. These cells also expressed the perivascular pre-FDC marker, platelet-derived growth factor receptor beta (PDGFRβ) (Fig. 6c, d). Therefore, marginal sinus FDC precursors expand during STm infection.

**Figure 6.**
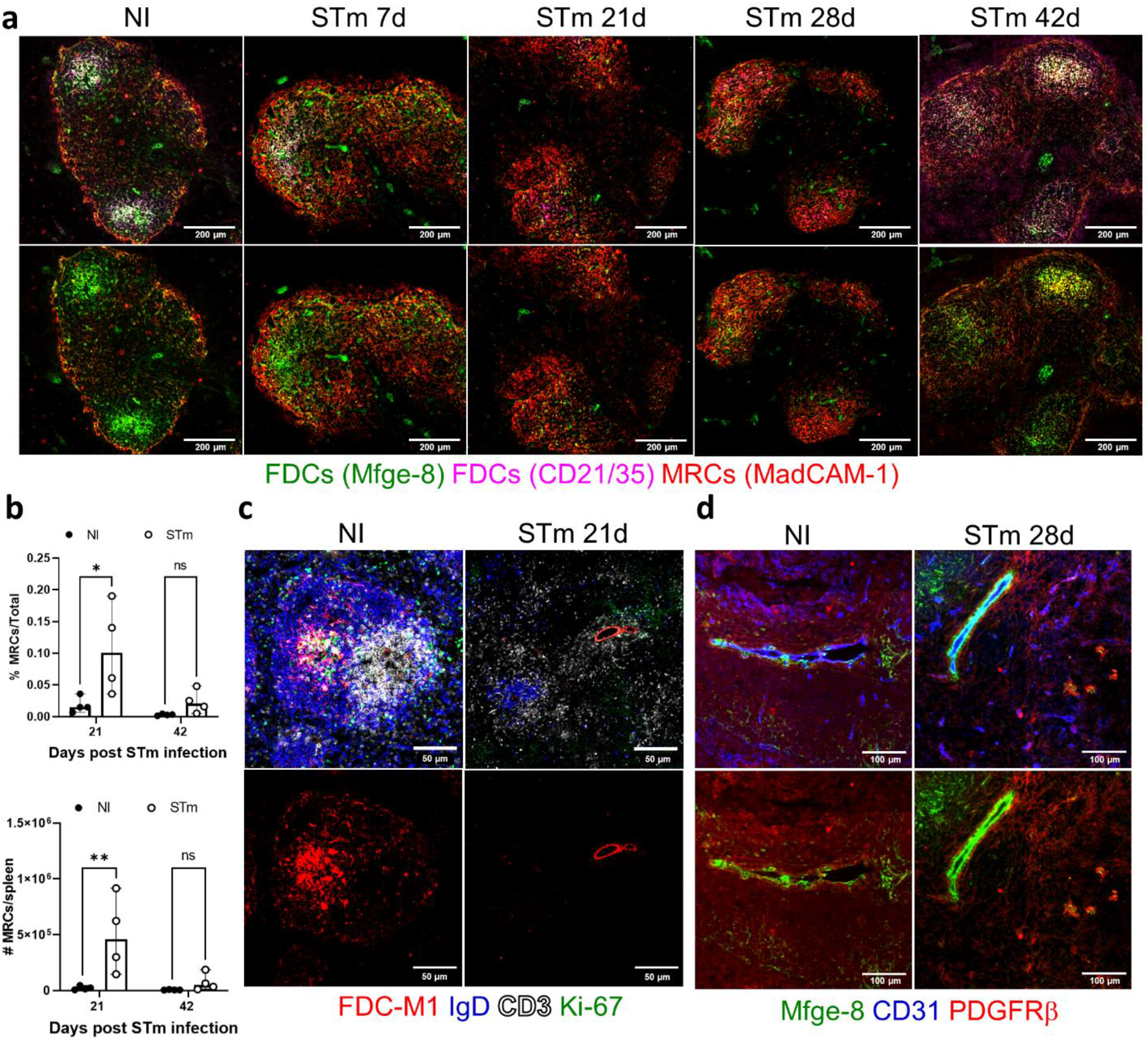
STm infection induces the expansion of MRCs. Mice were infected as per Fig. 1. **a** Representative IF images show MRCs (MadCAM-1+; red), FDCs (Mfge-8+; green), and CR1/2 (CD21/35; magenta). Top row images show all markers combined at different time points after infection; triple positive cells are present as white. The bottom row images show the merge of MadCAM-1 and Mfge-8 signal; double positive cells are present as yellow. Scale bar 200 μm. **b** Flow cytometry quantification of the proportion and absolute cell number of MRCs in NI mice and STm-infected mice after 21 days and 42 days. Cells were gated on CD45^-^CD31^-^TER119^-^podoplanin^+^MadCAM-1^+^CD21/35^-^. Each point represents one spleen, the bar height represents the median, and the error bars display the 95% CI. Two-way ANOVA and Šídák’s multiple comparison test between the NI and STm-infected group was performed. \**p* < 0.05, \*\**p* < 0.01; ns, nonsignificant. **c** IF images on top row show the merged staining for FDC-M1 (red), IgD (blue), CD3 (white) and Ki-67 (green). Bottom row displays single-colour images for FDC-M1 (red). Scale bar 50 μm. **d** Top panels show IF images for the combined staining of Mfge-8 (green), CD31 (blue) and PDGFRβ (red). Bottom row shows IF images for combined Mfge-8 and PDGFRβ staining. Scale bar 100 μm.

### MadCAM-1-expressing cells become major producers of CXCL13 during infection

Given the re-organisation of the WP and the lack of FDC networks induced after STm infection, we hypothesised that STm infection perturbs the expression of chemokine profiles within the WP. Key amongst these chemokines are CCL21 and CXCL13, which are required for normal T zone and follicle segregation. The distribution of these chemokines was examined by IF microscopy and gene expression from microdissected WP in spleens from non-infected mice and after 21 days of STm-infection. CCL21 gene expression was reduced in the WP and was also reflected at the protein level in T zones of infected mice compared to naïve controls (Fig. 7a, b; Supplementary Fig. 4a). In contrast, and despite perturbed FDC networks being detected, the expression of CXCL13 was comparable both when gene expression was measured in WP and protein expression in follicular areas (Fig. 7c, d; Supplementary Fig. 4b). In non-infected mice, most CXCL13 staining is associated with FDCs in follicles. In contrast, at day 21 post-infection, most CXCL13 is associated with MadCAM-1+ cells, which are distributed throughout the WP and are not restricted to the marginal sinus (Fig. 7e; Supplementary Fig. 4c). Thus, during STm infection MadCAM-1+ cells provide an alternative source of CXCL13 to compensate for the reduced expression of this chemokine by FDCs during this time.

**Figure 7.**
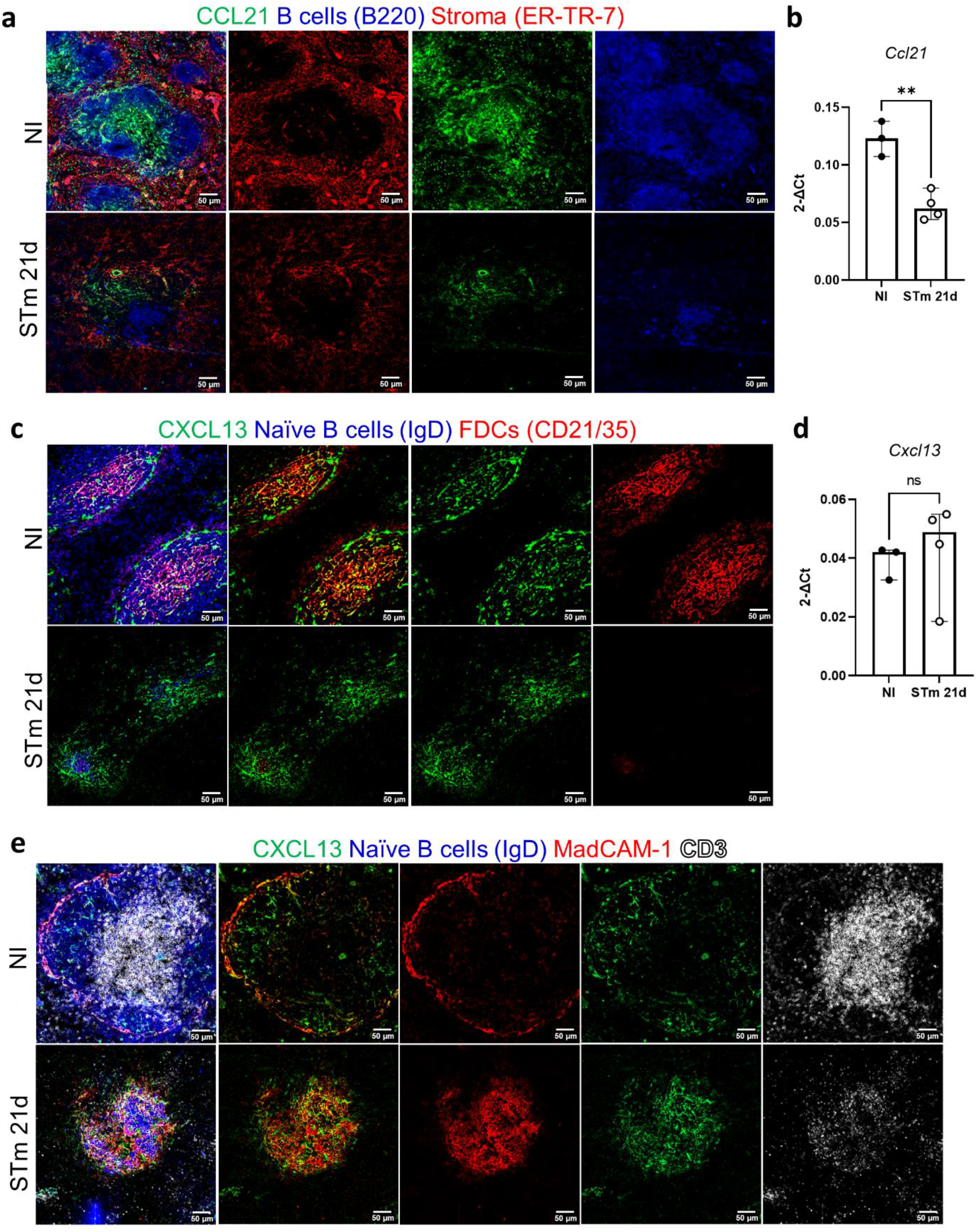
CXCL13 expression in the follicles is maintained after infection with STm. Mice were infected as per Fig. 1. **a** Representative IF images show CCL21 (green) expression along with B220 (blue) and ER-TR-7 (red) staining in NI mice (top row) and mice infected for 21 days (bottom row). Left-hand panels show merged images, and single-colour panels are displayed in the other panels. **b** Gene expression of *Ccl21* in WP of NI mice and mice infected with STm for 21 days. **c** Representative IF images of spleen sections stained for CXCL13 (green), IgD (blue), and CD21/35 (red) in control mice (top row) and STm-infected mice for 21 days (bottom row). Merged three-colour images (left), CXCL13 and CD21/35 merged images (second column), and single-colour channels (two columns to the right) are displayed. Scale bar 50 μm. **d** Graph represents gene expression of *Cxcl13* in WP from NI mice and mice infected with STm for 21 days. Each point in the graphs represents the gene expression detected in WP from one individual mouse, the bar height represents the median, and the error bars display the 95% CI. Two-tailed unpaired, *t*-test was used to compare groups. \*\**p* < 0.01, ns, nonsignificant. **e** Spleen sections were stained for CXCL13 (green), IgD (blue), MadCAM-1 (red) and CD3 (white). Representative IF images on top row show NI mice and images from STm-infected mice are shown in the bottom row. Images on the right displayed four-channel merged images, MadCAM-1 and CXCL13 merged images are presented in the second column and double-positive cells are yellow. Single-colour images are shown to the left. Scale bar 50 μm.

## Discussion

Here, we show how systemic *Salmonella* infection induces the disorganisation of the splenic WP and loss of GC formation, and the later detection of GCs correlates with the induction and resolution of these changes. The capacity of STm infection to impact GC induction and maintenance is not restricted to STm-specific responses but also impacts GC induced to a diverse spectrum of other pathogens and antigens, including model antigens, bacterial flagellin, the microbiota, influenza virus and the helminth *Nippostrongylus brasiliensis*^25–28^. The diversity of these antigens and the observation that STm infection impacts pre-existing and ongoing GC responses indicates the antigen independence of these effects. Since B cells from STm-infected mice maintain the capacity to develop into GC B cells^28^, it indicates that a significant contributory reason for the absence of GC in the first weeks of infection is that STm disrupts the niches in which GCs develop.

Previous studies have shown how non-B cell-intrinsic mechanisms including the recruitment of discrete Sca-1+ monocyte populations, TLR4 expression and IL-12-mediated suppression of Tfh cells can impair GC formation after STm infection^27,28^. The impact of IL-12 induced to *Salmonella* is likely to act early, possibly in the first 24 hours after infection since this cytokine is produced by conventional and monocyte-derived dendritic cells from 2 hours post-infection to promote Th1 differentiation^26^. In contrast to these early events, the identification of perturbed FDC and MZ organisation occurs later and indicates that the structures needed to support GC development and function are not maintained during the peak of the infection. Despite the multiple and quite marked effects of *Salmonella* on the host adaptive immune response, it is important to contextualise the lack of GCs as being a selective effect and not representative of a general impairment in the capacity of the host to induce antibody responses to the pathogen. Many B cell responses remain active in the host during the period when GC are not detectable. For instance, extensive EF B cell responses are detectable from the first days after infection^20^. These responses are induced in a T-independent manner with B cell switching to IgG dependent upon BCL6^+^PD1^lo^CXCR5^lo^ T cells and CD40L^20,38,39^. Moreover, the EF response-derived antibody is functional and can moderate bacteraemia and subsequent re-infection^20^. Therefore, the tissue reorganisation observed in primary infection is not a barrier to productive EF responses and so the impairment of the GC response to *Salmonella* is unlikely to have evolved simply as a strategy to impair antibody responses *per se*. Therefore, whether there is a selective advantage to the host or to the pathogen remains unclear.

A hallmark of the GC is the organisation of multiple cell types within follicles. This includes FDCs which are essential not only for providing chemokines to recruit B cells but also for antigen-driven selection^12,16,40^. The STm-induced changes in FDCs and the downregulation of key phenotypic markers, alongside the timing of these changes, are fully consistent with changes in FDCs being a key reason why GC do not form or are not maintained. Nevertheless, it is unlikely that direct infection of FDCs drives these effects as bacteria are rarely detected in FDC and B cell follicles, but mainly localise in iNOS+ inflammatory foci in the red pulp during the weeks after infection examined here^41,42^. This contrasts with what is observed after other infections where impaired GC responses associated with disruption of FDCs have been observed. For instance, FDCs are directly infected and lost subsequently in draining LNs after the Bluetongue virus infection of sheep^43^.

FDCs are plastic stromal cells that originate from perivascular cells and require B cell-derived TNFR and LTβR signalling for their maintenance^35,37,44–47^. The de-clustering of FDC induced by STm infection is unlikely to reflect the death of FDCs as although the density of FDCs in the spleen was significantly reduced after infection, the total numbers were not and these cells are known to be long-lived and resistant to stresses such as radiation exposure^15,48^. Another potential outcome is that FDCs lose maturity and partially de-differentiate as FDC-like cells were readily detectable in the follicles and expansion of MadCAM-1+ MRCs in the MZ and throughout the follicle was observed. Likewise, the capacity to form normal FDC networks could also occur due to the reduced density of B cells in the follicles or/and an intrinsic decrease of LTαβ expression, which may limit LTβR signalling. The other LTβR ligand, LIGHT (*TNFSF14*) may contribute to this process^49^. Here, we have shown that the expression of genes within the TNFR and LTβR signalling pathways in the WP were modulated by STm infection. TNF is produced by multiple innate immune cell types and T cells after infection and plays pleiotropic roles in controlling infection in tissues such as the spleen and liver^50–52^. Less is known about the contribution of lymphotoxin and TNF to the organisation of WP after STm infection. We did attempt to modulate FDCs by targeting the LTβR and modulation of TNF using agonistic and neutralising antibodies, respectively. Targeting LTβR did not provide consistent results, with some mice showing accelerated induction of FDCs and GCs, but not all. The reasons for this are unclear but we speculate that they may be consequences of trying to maintain agonistic LTβR signalling for a sufficiently long time to have lasting effects. In contrast, neutralising TNF during *Ehrlichia muris* infection enhances GC responses^21^ but we observed an enhanced loss of FDCs and no GC response during STm-infection. The loss of FDCs, GCs and impaired humoral immunity after interference with the TNF signalling using either TNF-neutralising antibodies or gene-targeted mice has been described extensively in steady-state conditions^36,46,47,53,54^ and under inflammatory environments such as sepsis^55^. These results suggest that during active infection, the targeting of specific cell types may be more difficult than targeting soluble factors such as TNF and highlights the potential difficulty in using single agent interventions to target different bacterial infections as each infection may have a distinct immune signature.

## Acknowledgements

This work was supported by the Royal Society under the Newton International scheme to E.M.J and A.F.C (NIF\R1\192061) and by Medical Research Council (MR/N023706/1) grant to A.F.C. AA, RP, SJ and RL are PhD students supported by King Saud University, Riyadh, South Arabia, Wellcome AAMR DTP scholarship, BBSRC, and Wellcome Trust (222389/Z/21/Z, part of 108871/B/15/Z), respectively. The funders had no role in study design, data collection and interpretation, or the decision to submit the work for publication. We thank the Microscopy and Flow Cytometry Services, and staff within the Biomedical Services Unit at the University of Birmingham for their technical help with experiments. Especially we thank Professor Carl F. Ware (Sanford Burnham Prebys Institute) for his insightful comments and feedback on the manuscript.

## Author Contribution

EMJ, AFC, and FB conceived and designed the experiments. EMJ, MPT, AA, RP, SJ, and RL performed the experiments. EMJ, SN, and EP performed the laser capture microdissection and RNA data analysis. EMJ, SN, EP, and AFC analysed the data. EMJ, AFC, MPT, SN, IRH, FB, and AFC interpreted and discussed the data. EMJ and AFC wrote and revised the manuscript. All authors commented on drafts of the manuscript and approved the final version.

## Supplemental information titles and legends

**Supplementary Figure 1.**
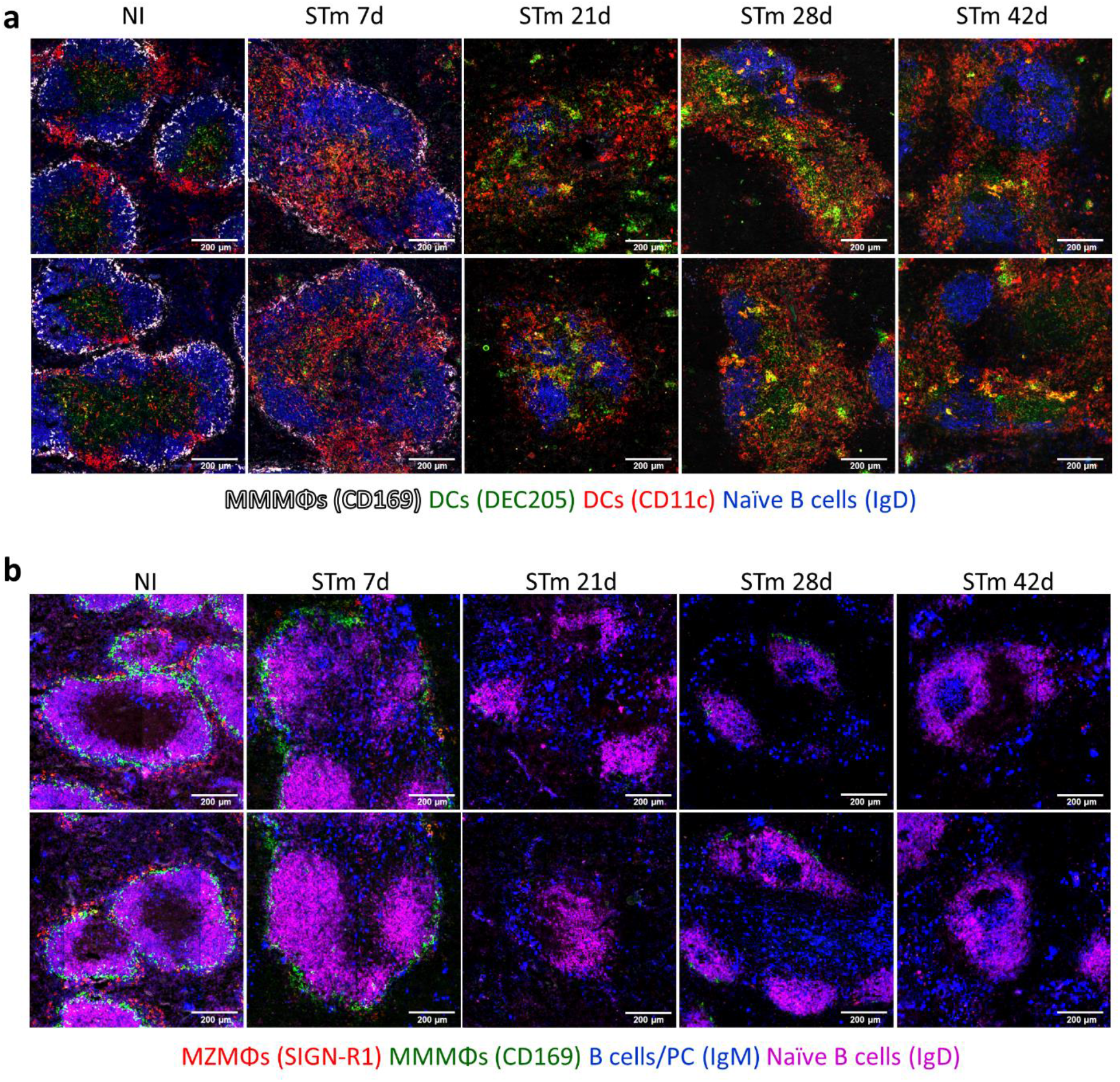
DCs, MZMΦs, MMMΦs and MZ B cells in the spleen after STm infection. Mice were infected as per Fig. 1. **a** Cryosections from spleens were stained to detect MMMΦs (CD169+; white), DCs (DEC205+; (green) and CD11c+ (red)), and naïve B cells (IgD+; blue). Scale bar 200 μm. **b** Cryosections from spleens were stained to detect MMMΦs (CD169+; green), MZMΦs (SIGN-R1+; red), MZ B cells (IgM+ cells in the MZ; blue) and B cells (IgD+; magenta). Scale bar 200 μm.

**Supplementary Figure 2.**
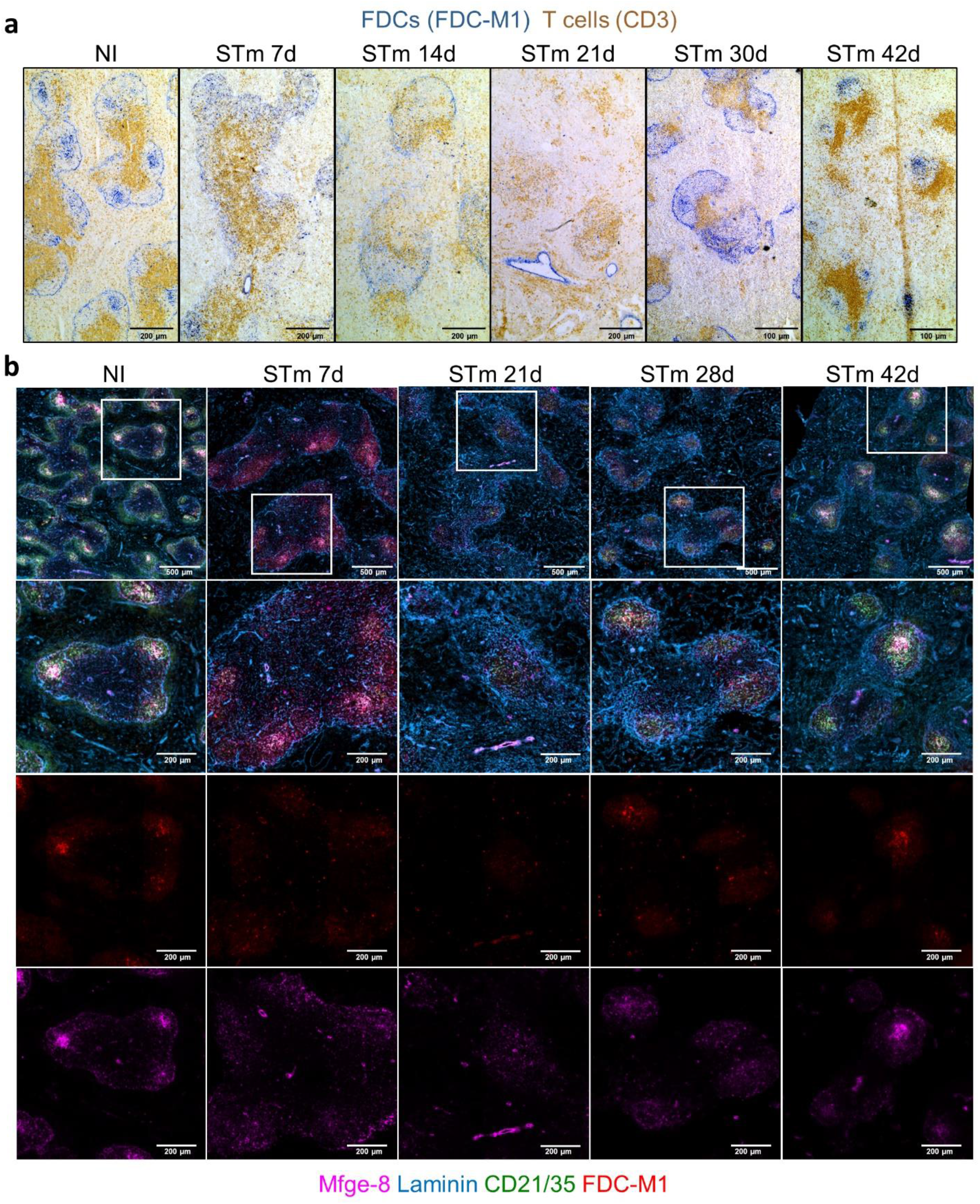
FDC analysis in the spleen during STm infection. Mice were infected as per Fig. 1. **a** Representative images of spleen cryosections stained by immunohistochemistry to detect FDC (FDC-M1+; blue) and T cells (CD3+; brown). Scale bar 200 μm. **b** IF images stained to detect laminin (blue), FDCs (Mfge-8+; magenta and FDC-M1; red), and CR1/CR2 (CD21/35+; green). Top row shows low magnification and merged images of all markers, and the second row represents higher magnifications of the selected areas. The bottom two rows show single-colour images. Scale bar 200 μm.

**Supplementary Figure 3.**
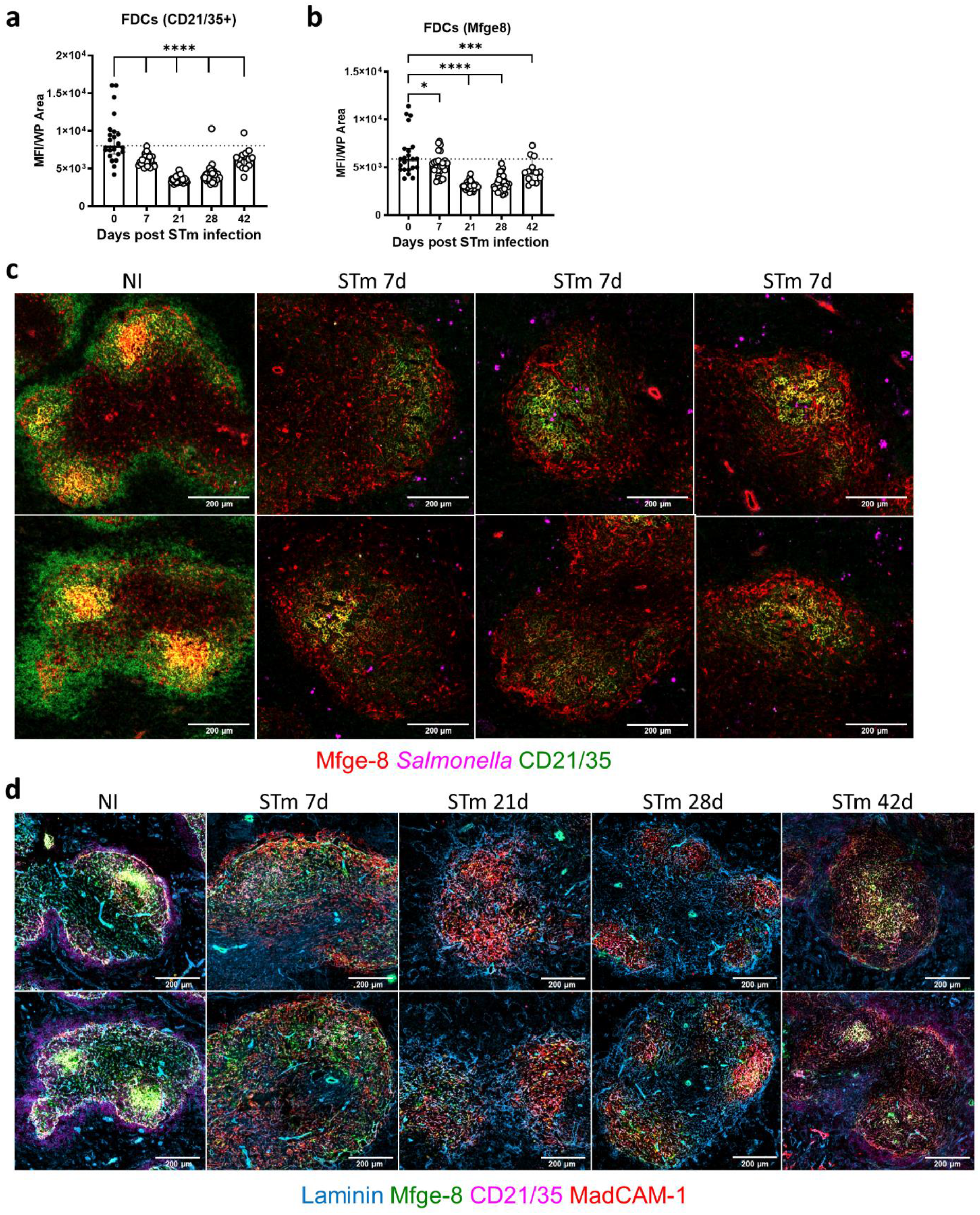
Distribution of STm and FDCs in the WP of the spleen 7 days after infection. Mice were infected as per Fig. 1. **a,b** Graphs display the *in situ* quantification of the MFI for the detection of CD21/35 and Mfge-8 at different time points after STm infection in individual WP. Each symbol represents the MFI per WP, the bar height represents the median, and the error bars display the 95% CI. One-way ANOVA and Dunnett’s multiple comparison test was performed for **a-b**. \**p* < 0.05, \*\*\**p* < 0.001, \*\*\*\**p* < 0.0001. **c** Cryosections were stained to detect FDCs (Mfge-8+; red), CR1/2 (CD21/35+; green) and STm (magenta) in NI mice (left column) and mice infected for 7 days (six images to the right). The yellow signal represents Mfge-8 and CD21/35 double positive cells. Scale bar 200 μm. **d** Representative IF images show MRCs (MadCAM-1+; red), FDCs (Mfge-8+; green), CR1/2 (CD21/35; magenta), and laminin (blue). Scale bar 200 μm.

**Supplementary Figure 4.**
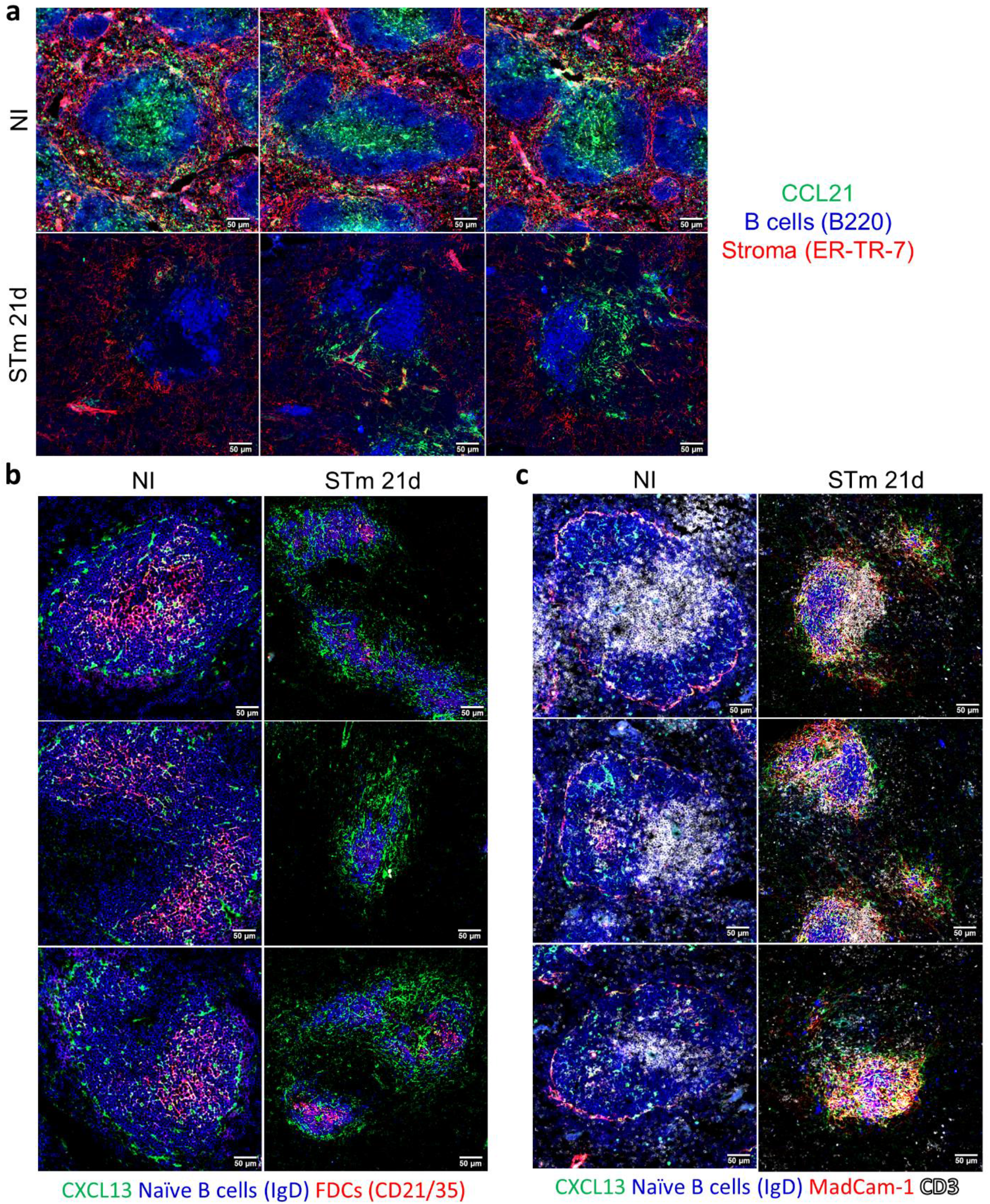
Chemokine expression in the splenic WP after STm infection. Mice were infected as per Fig. 1. **a** Representative IF images show CCL21 (green) expression along with B220 (blue) and ER-TR-7 (red) staining in NI mice (top row) and mice infected for 21 days (bottom row). Scale bar 50 μm. **b** Representative IF images of spleen sections stained for CXCL13 (green), IgD (blue), and CD21/35 (red) in control mice (left) and mice infected with STm for 21 days (right). Scale bar 50 μm. **c** Spleen sections from NI mice (left) and STm-infected mice (right) were stained for CXCL13 (green), IgD (blue), MadCAM-1 (red) and CD3 (white). Scale bar 50 μm.

